# High-throughput Raman-activated cell sorting in the fingerprint region

**DOI:** 10.1101/2021.05.16.444384

**Authors:** Matthew Lindley, Julia Gala de Pablo, Jorgen Walker Peterson, Akihiro Isozaki, Kotaro Hiramatsu, Keisuke Goda

## Abstract

Cell sorting is the workhorse of biological research and medicine. Cell sorters are commonly used to sort heterogeneous cell populations based on their intrinsic features. Raman-activated cell sorting (RACS) has recently received considerable interest by virtue of its ability to discriminate cells by their intracellular chemical content, in a label-free manner. However, broad deployment of RACS beyond lab-based demonstrations is hindered by a fundamental trade-off between throughput and measurement bandwidth (i.e., cellular information content). Here we overcome this trade-off and demonstrate broadband RACS in the fingerprint region (300 − 1,600 cm^−1^) with a record high throughput of ~50 cells per second. This represents a 100× throughput increase compared to previous demonstrations of broadband fingerprint-region RACS. To show the utility of our RACS, we demonstrate real-time label-free sorting of microalgal cells based on their accumulation of carotenoids and polysaccharide granules. These results hold promise for medical, biofuel, and bioplastic applications.

## Introduction

Cell sorting is the workhorse of biological research and medicine^1^. Cell sorters are commonly used to sort heterogeneous populations of cells based on their intrinsic features and have numerous applications in microbiology, immunology, virology, stem cell biology, and metabolic engineering^2–6^. While several schemes have been developed for sorting cells based on features such as size^7^, morphology^8,9^, and surface antigens^10^, those able to sort at high throughput for intracellular chemical content are limited to fluorescence-activated and Raman-activated cell sorting (FACS and RACS, respectively)^1,11^. In both methods, sorting is accomplished by a stepwise automatic process: 1. measurement of chemical content with single-cell resolution, 2. analysis of the measurement data to identify cells of interest, and 3. mechanical separation of target cells from the original sample^1^. FACS machines commonly identify molecular targets by signals from attached exogenous fluorophores or co-expressed transgenic fluorescent proteins. This indirect detection scheme is often referred to as labeled detection^1^. Although fluorescent labeling is essential for microbiological research, the use of labels has significant drawbacks. These include reduced cellular vitality due to the cytotoxicity of labels themselves, nonspecific binding of labels, a lack of labels or transfection methods for many molecular targets and species, incompatibility with human-targeted cell therapies, and increased experimental complexity imposed by labeling protocols^11^. On the other hand, RACS machines directly identify target molecules in a label-free manner via Raman spectroscopy, which measures the inelastic scattering of incident photons by characteristic molecular vibrations^11^. To date, FACS technology is well developed, commercialized, and broadly adopted, while RACS is nascent and comprised of a small number of lab-based demonstrations^11,12^.

The fundamental downside of RACS, which hinders its broad deployment in biomedical fields, is a trade-off between throughput and measurement bandwidth (i.e., cellular information content) (Fig. 1). Since Raman scattering is a low-probability phenomenon, broadband spontaneous Raman spectroscopy requires long signal integration times, typically on the order of >100 ms. This has resulted in schemes where RACS machines use laser tweezers to catch cells from the flow stream and hold them for measurement^13^, or laser ejection schemes where slide-affixed cells of interest are measured and subsequently ejected from their substrate by an intense laser pulse^14^. Although these catch-and-release-based schemes produce broadband informative single-cell Raman spectra, they suffer from throughputs of <0.1 events per second (eps)^14–21^, where an event is defined as the triggering of a measurement by the passage of a cell, cell cluster, or cell-like debris. To date, flow-through broadband spontaneous RACS machines demonstrate throughputs below ~0.5 eps at best^22^. To overcome this, a higher throughput alternative is offered by coherent Raman spectroscopy, which utilizes intense laser pulses to drive coherence among the vibrational modes of the sample molecules. These coherences can produce signal improvements of ~5 orders of magnitude^23^, facilitating rapid spectral acquisition (sub ms) and thereby higher throughput. This was recently shown by Nitta *et al*., with a markedly higher cell sorting throughput of 46 eps via stimulated Raman scattering (SRS) measurement. However, this method confined detection to the CH-stretching region (2,800 – 3,100 cm^−1^) and measured only four spectral points^12^. Thus, current RACS demonstrations leave a significant gap between the low-throughput spectrally rich spontaneous fingerprint techniques and the high-throughput but spectrally limited SRS-based techniques.

**Fig. 1.**
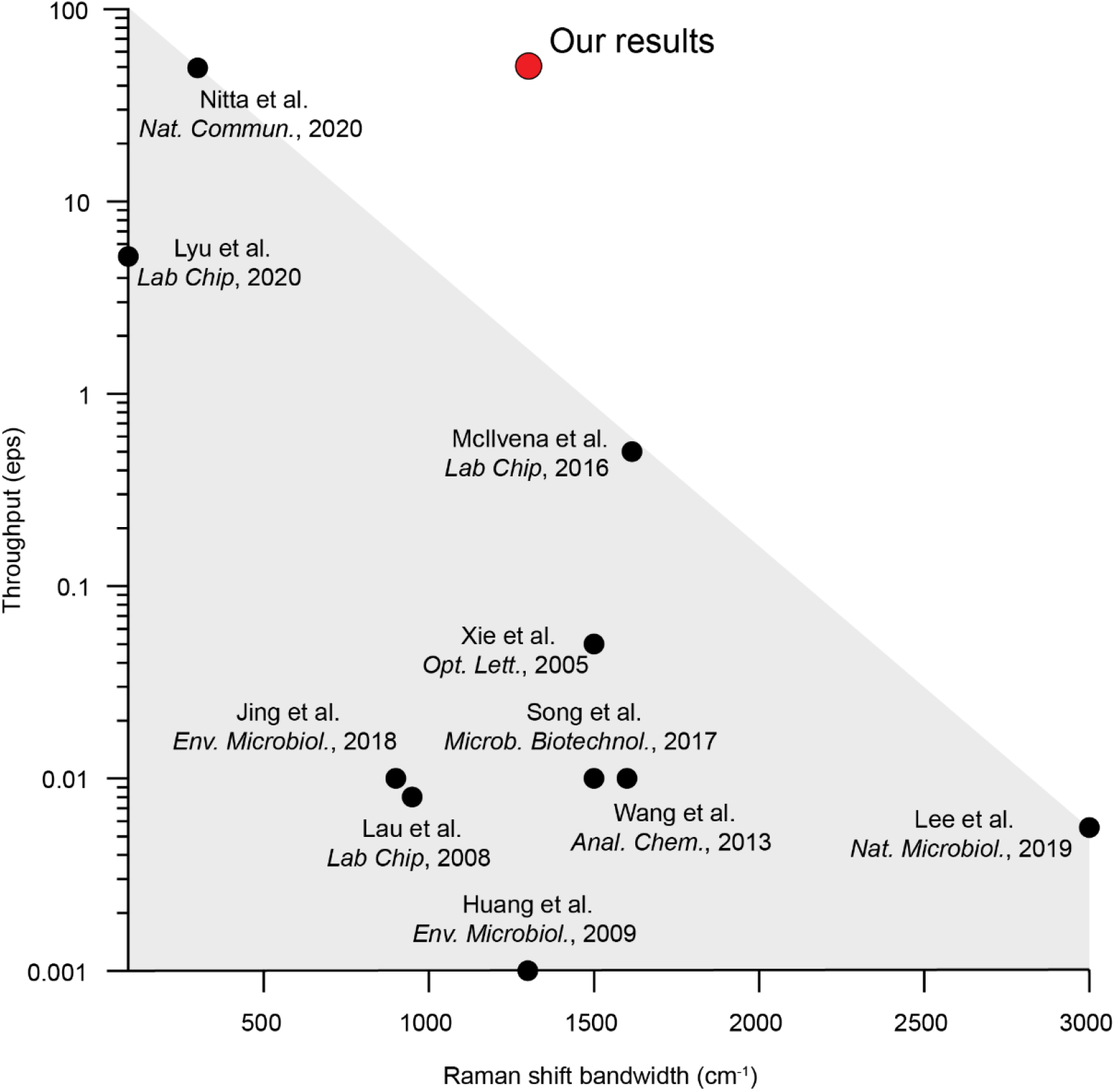
Trade-off between throughput and measurement bandwidth (i.e., cellular information content) in RACS demonstrations. RACS demonstrations to date leave a significant gap between the spectrally rich but low-throughput spontaneous fingerprint techniques and the high-throughput but spectrally limited SRS-based techniques. Our RACS machine overcomes the trade-off and demonstrates a 100× improvement in throughput compared to previous RACS demonstrations at similar bandwidths.

In this Article, we break this trade-off by combining a rapid-scan Fourier-transform coherent anti-Stokes Raman scattering (FT-CARS) spectrometer^24–26^, an on-chip dual-membrane push-pull cell sorter^27^, and a real-time spectral processor built on a field-programmable gate array (FPGA), and demonstrate broadband RACS in the fingerprint region (300 – 1,600 cm^−1^) with a high throughput of up to 50 eps. This represents a 100× throughput increase compared to previous broadband fingerprint-region RACS demonstrations (Fig. 1), allowing our RACS machine to finish in a few minutes experiments that would take previous broadband RACS machines several hours. This offers a significant advantage for large-scale and time-sensitive cell sorting. We typify sorting performance with polymer microbeads, achieving sorted purities as high as 98%. To show the utility of our RACS machine, we demonstrate real-time label-free sorting of microalgal cells based on their intracellular accumulation of polysaccharide granules and carotenoids. This demonstration also distinguishes RACS and FACS; while microalgal polysaccharide granules continue to garner interest for medical^28^, biofuel^29,30^, and bioplastic^31^ applications, they remain difficult to stain and thus detect in FACS machines, highlighting the suitability of our RACS machine for unmet bioengineering applications.

## Results

As shown in Fig. 2, our RACS machine combines a FT-CARS spectrometer, an on-chip dual-membrane push-pull cell sorter, and a real-time spectral processor that provides spectral analysis and sort signaling. The spectrometer, described in our previous work^24^, rapidly scans the optical delay between a pair of femtosecond pulses, with the first pulse driving a vibrational coherence in the sample and the second pulse probing the coherence across increasing delay. The resulting anti-Stokes scattering intensity is modulated across this delay by the time evolution of sample vibrations, allowing Raman spectra to be recovered by Fourier transform. For this work, the spectrometer was tuned to measure 24,000 spectra per second, with the spectral region of 300 – 1,600 cm^−1^ and a spectral resolution of 30 cm^−1^. Cells in suspension were injected into the microfluidic chip and focused into a single stream within the main channel (200 μm × 200 μm cross-section) using an on-chip acoustic focusing piezoelectric transducer. The chip position was adjusted so that the cells passed through two successive measurement regions. In the first region, a cell’s passage was detected by the forward-scattering of a 632-nm diode laser, to produce a trigger signal. In the second region, FT-CARS spectra were measured in response to the trigger. The forward scattering and FT-CARS focal positions were held static throughout the measurement. We calculated the FT-CARS focal diameter as 0.5 μm based on the beam diameter and wavelength of the laser light and the numerical aperture of the objective lens. Upon event triggering, raw measurement data was passed to the FPGA, which produced and averaged the first 2^N^ spectral points (where N is a user-selected integer from 0 to 5) to make a representative spectrum for the event. Using this spectrum, the FPGA then assigned a sort decision based on analysis of spectral features. After the measurement region, the microfluidic channel split into three outlet channels for binary (sort/unsort) sorting. Under normal flow conditions and without a sort decision, cells passaged down the central channel to the unsort collection. For a sort decision, the dual-membrane push-pull sorter generated a jet of water across the main channel, pushing the target cell into either of the side channels, which joined downstream for collection. This allowed selected cells to be sorted by both push and pull actuation, maximizing the sorter duty cycle^27^.

**Fig. 2.**
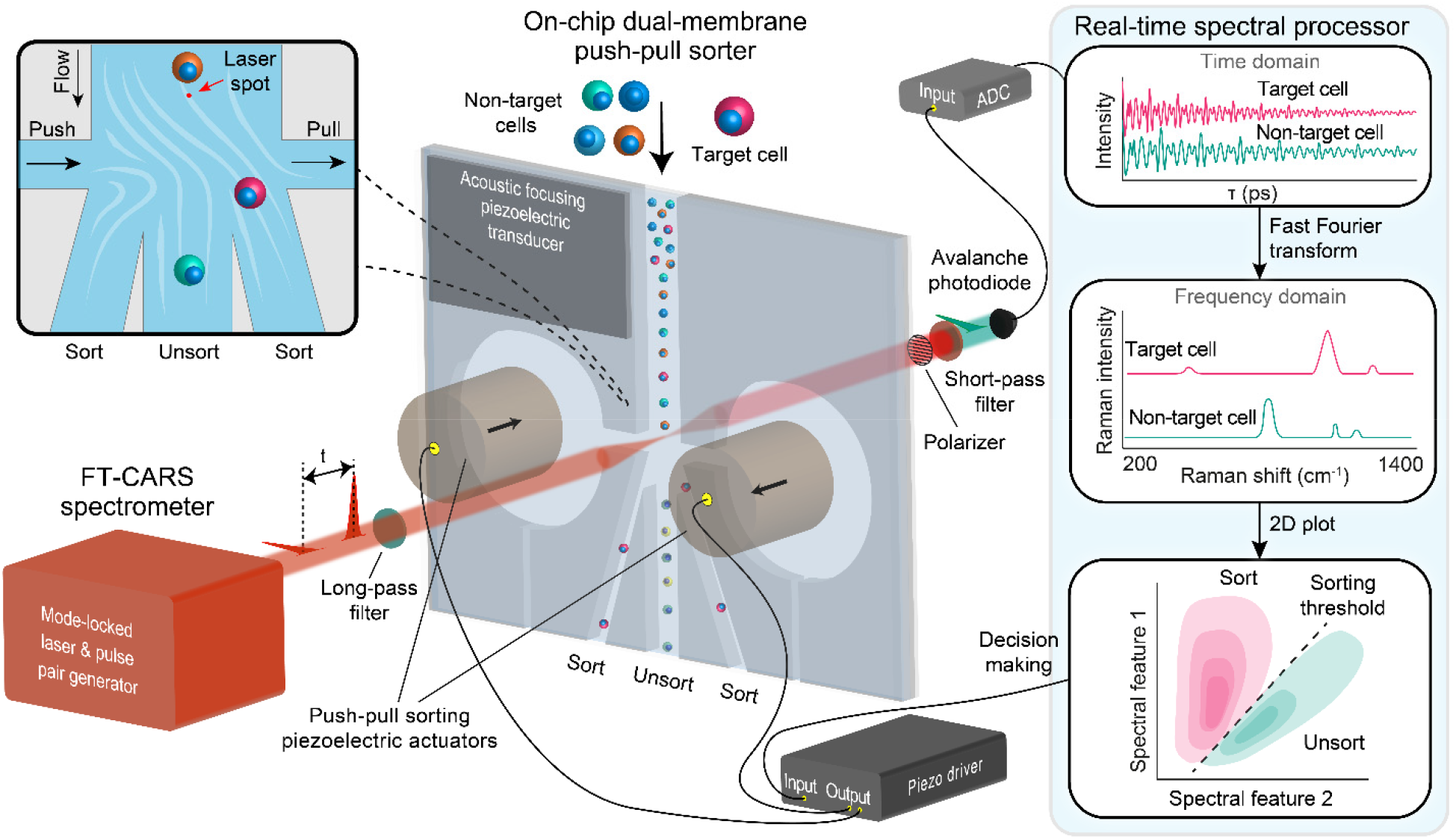
Schematic of the high-throughput, broadband RACS machine. The RACS machine combines an FT-CARS spectrometer with an on-chip dual-membrane push-pull sorter and a real-time spectral processor. Cells in suspension are injected into the microfluidic channel of the RACS machine and focused into a single stream by the acoustic-focusing piezoelectric transducer as they flow. Cells are then measured as they pass though the FT-CARS beam focus. Raman scattering data are recorded by the avalanche photodiode as a time-domain interferogram and passed to the real-time spectral processor for automated analysis (**right inset**). The time-domain waveform is Fourier-transformed to produce a broadband fingerprint-region Raman spectrum for each cell. Spectral features are analyzed to make a “sort” or “unsort” decision for the cell. For a sort decision, the dual-membrane push-pull sorter generates a jet of water across the channel, carrying the cell out of the main flow channel and into a side channel for sort collection (**top left inset**). The push-pull actuation is not undertaken for cells receiving an unsort decision, allowing them to continue along the main channel and into the unsort collection. ADC, analog-to-digital converter.

We typified the sorting performance of our RACS machine by sorting a 1:1 mixture of poly(methyl methacrylate) (PMMA, 5 μm in diameter) and polystyrene (PS, 7 μm in diameter) microbeads based on the intensity ratio of their Raman peaks at 810 cm^−1^ and 1,003 cm^−1^ (A microbead sorting video is available as Supplementary Movie 1). Two trials were performed in which the samples were sorted at throughputs of 31 and 13 eps. Averaged spectra from the measurement of the microbeads (2,892 for PS and 10,293 for PMMA) are shown in Fig. 3A, identifying typical peaks of PS and PMMA. Scatter plots of per-microbead peak intensities are shown in Figs. 3B, with the sorting thresholds indicated by the black curves. The microbeads were fluorescent-labeled, allowing the sorting results to be quantified via fluorescence microscopy as shown in Fig. 3C. Quantification results are shown in Fig. 3D. Our RACS machine achieved 98% purity and 69% yield for PS selection at 13 eps and 90% purity and 53% yield at 31 eps. Assuming ideal flow focusing, measurement, and sorting, the fundamental limit to purity is determined by the probability that an event contains only one particle or cell^12^. Our results for microbead sorting are compared to this model in Fig. 3E, which shows that our RACS machine achieved the theoretical maximum sorting purity for microbead sorting at 13 eps and was 4.0% below maximum at 31 eps.

**Fig. 3.**
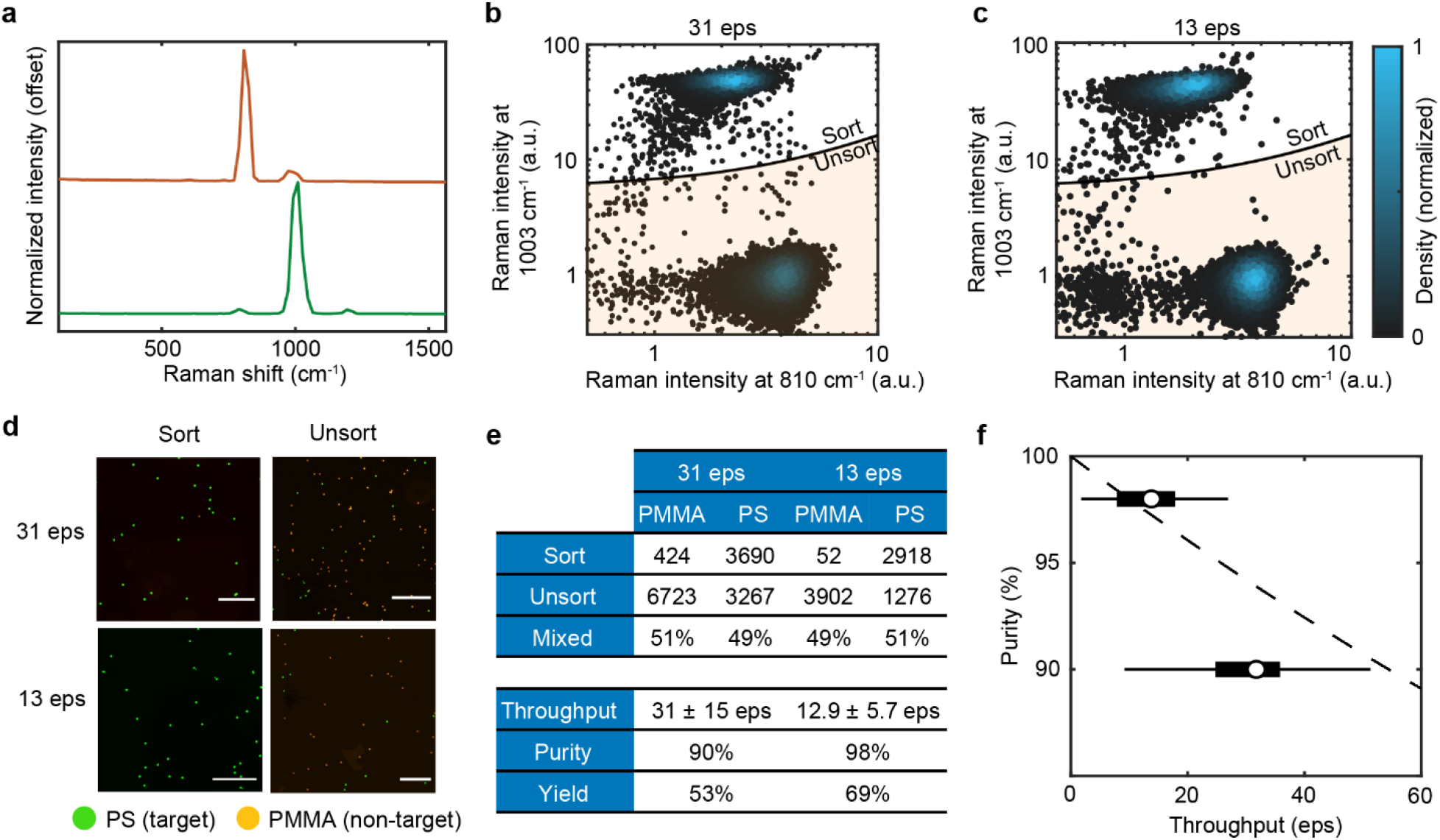
Characterization of sorting performance with microbeads. (**a**) Averaged spectra from PS and PMMA microbeads under flow at 31 eps. (**b**) Scatter plots of per-microbead peak intensities for sorting at 31 eps and (**c**) 13 eps. The sorting threshold is indicated by the black curve. (**d**) Fluorescence images of collected microbeads from the sort and unsort outlets. PS is green and PMMA yellow. (**e**) Summary of sorting results. (**f**) Resulting purities (dots) from 31 eps and 13 eps sorting with throughput distributions shown as a box plot. The theoretical maximum of purity vs throughput for a 1:1 mixture is included (dashed line).

To show the broad utility of our RACS machine, we sorted cell samples from the microalga *Euglena gracilis* (NIES-48) based on paramylon content, *Chromochloris zofingiensis* (NIES-2175) based on starch content, and a mixed sample of *Haematococcus lacustris* (NIES-4141) and *E. gracilis* based on astaxanthin content. These three experiments demonstrated the RACS machine’s sensitivity for polysaccharides and carotenoids and, in the latter two experiments, showcased the RACS machine’s sorting performance when handling samples of mixed cell sizes and morphological features. Bright-field images of each species passing from the measurement region to the sorting region in the on-chip dual-membrane push-pull cell sorter are shown in Figs. 4A-4C. Corresponding single-cell and ensemble average spectra taken by the RACS machine during measurement for these species are shown in Figs. 4D-4F. Averaged spectra from standard samples of paramylon, starch, and astaxanthin were included for comparison. The peaks in the cell spectra were in agreement with the standard samples. The FT-CARS laser power was 150 mW at the beam focus for paramylon- and starch-targeted sorting. Astaxanthin was measured in the resonance Raman regime with a reduced laser power of 94 mW.

**Fig. 4.**
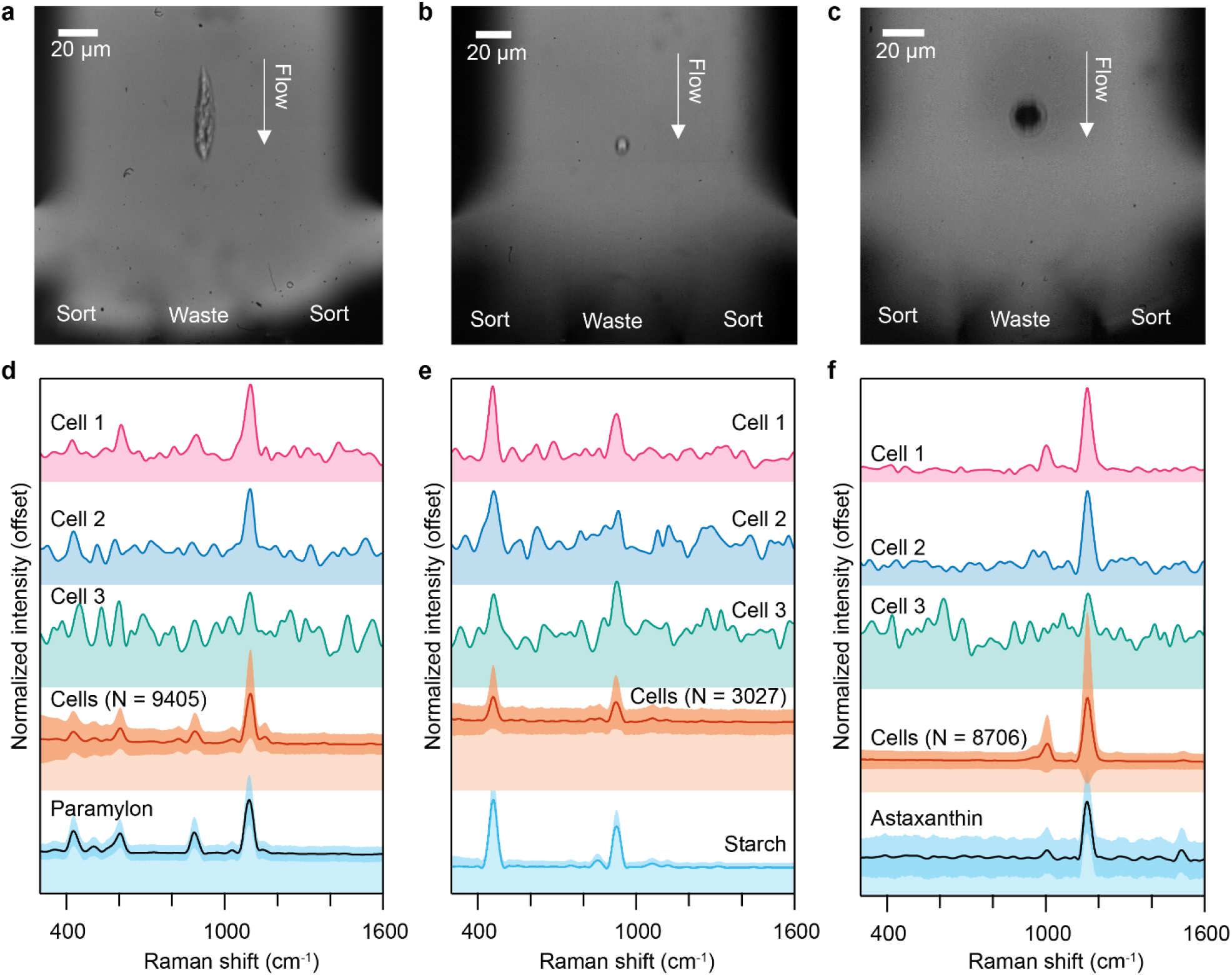
Cell images and spectra from the high-throughput, broadband RACS machine. Bright-field images of (**a**) an *E. gracilis* cell, (**b**) a *C. zofingiensis* cell, and (**c**) a *H. lacustris* cell passing from the measurement region to the sorting region in the on-chip dual-membrane push-pull sorter. Also shown are single-cell and ensemble average spectra taken by the RACS machine during measurement of (**d**) *E. gracilis* cells, (**e**) *C. zofingiensis* cells, and (**f**) *H. lacustris* cells. Averaged spectra form standard samples of paramylon granules, corn starch granules, and astaxanthin included for comparison. All spectra shown in the figure are normalized for peak height at the maximum peak position of the standard samples. Darker shading along averaged spectra indicates ±1 standard deviation.

We first sorted *E. gracilis* cells for paramylon content. *E. gracilis* is a free-swimming, single-cell, mixotrophic microalga that accumulates up to 80% (w/w) of its dry weight as granules of the β-1,3 glucan paramylon^32^. Paramylon and its derivatives have demonstrated immunomodulating^33,34^, anti-cancer^35^, and anti-HIV properties^36^ among other biomedical effects^37^. Additionally, paramylon has garnered interest as a precursor for biofuel^38^ and bioplastic^39^ production. For sorting, separate samples of *E. gracilis* cells were cultured under fully heterotrophic (dark) and mixotrophic (light) conditions, with the former heavily promoting paramylon accumulation and the latter suppressing it^40^. The presence of chlorophyll in light-grown cells and its absence in the dark-grown cells allowed chlorophyll autofluorescence to distinguish the samples when imaging the sorted results. Cells from each culture were mixed at a 1:1 ratio and sorted to select for paramylon-rich cells (determined by the Raman peak intensity at 1,092 cm^−1^), which were expected to originate from the dark-grown sample. The scatter plot of sorting results is shown in Fig. 5A, with an image of cells in the sort and unsort collections and a chart of the counted results shown in Fig. 5B. A total of 11,831 cells were sorted at a throughput of 47 eps, with a resulting purity of 92.7% and yield of 18.7%. Additional images of the sorting results are shown in Supplementary Fig. 1.

**Fig. 5.**
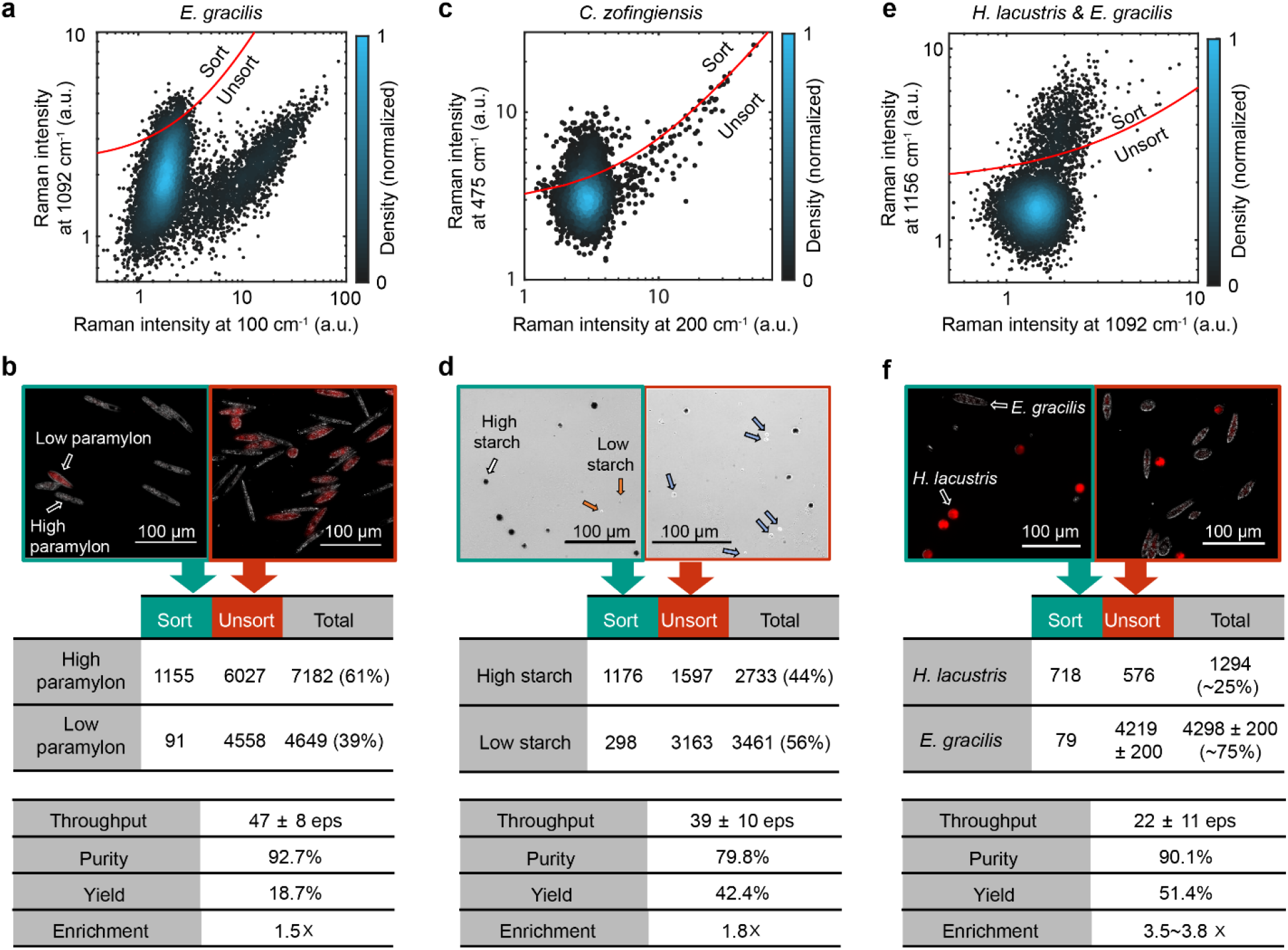
Demonstration of RACS. (**a**) Scatter plot from *E. gracilis* mixed phenotype sorting to select paramylon-rich cells, with (**b**) images from sort and unsort collections and counted results. (**c**) Scatter plot from mixed phenotype sorting of *C. zofingiensis* to select starch-rich cells, with (**d**) images from sort and unsort channels and counted results. (**e**) Scatter plot from sorting a mixed cell sample of *H. lacustris* and *E. gracilis* to select astaxanthin-rich cells, with (**f**) images from sort and unsort channels and counted results. Cell images in **b** And **f** Consist of overlaid bright-field (white) and chlorophyll fluorescence (red) intensities. Cell images in **d** Consist of bright-field intensities only.

To show sorting of small-sized (2-10 μm) cells, we next sorted *C. zofingiensis* cells for starch content. *C. zofingiensis* is a single-cell, mixotrophic microalga that accumulates starch when grown in media with high glucose concentration^41^. *C. zofingiensis* (previously called *Muriella zofingiensis* or *Chlorella zofingiensis*), sorted here, has been a focus of bioproduction discussion due to its high biomass yield under a wide range of culture conditions^42^. *C. zofingiensis* samples were prepared separately in high-glucose (modified Acetate with 20g/L glucose) and low-glucose (0.4 g/L glucose) media. The samples were then mixed at a 1:1 ratio and sorted based on their starch signal, determined by the FT-CARS peak intensity at 475 cm^−1^. The scatter plot of sorting results is shown in Fig. 5C, with images of cells in the sort and unsort collections and a chart of the counted results shown in Fig. 5D. A total of 6,234 cells were sorted at an average throughput of 39 eps. The phenotype of the cells was checked by Lugol’s iodine staining, which stained the starch-rich cells in the sort and unsort collections dark brown. Counting the results yielded a sorting purity of 79%, a yield of 42.4%, and an enrichment of 1.63×. A second sample mixed at a 1:9 ratio of the high:low glucose media cohorts was sorted to select starch-rich cells, resulting in a purity of 33.4%, yield of 45.2%, and enrichment of 6.9×, as shown in Supplementary Fig. 2. Additional images of the sorting results are shown in Supplementary Fig. 3 and a video of *C. zofingiensis* sorting is available as Supplementary Movie 2.

Finally, to show sorting performance with a mixed species cell sample, we sorted a mixed sample of *H. lacustris* and *E. gracilis* by their carotenoid content. *H. lacustris* accumulates the carotenoid astaxanthin when grown under stressful conditions such as nutrient deprivation^43^. In this sorting demonstration, astaxanthin-rich *H. lacustris* cells were separated from the mixed sample based on the intensity of the resonance Raman astaxanthin peak at 1,156 cm^−1^. The scatter plot of sorting results is shown in Fig. 5E, with images of cells in the sort and unsort collections and a chart counted results shown in Fig. 5F. The sorting performance was evaluated by identifying spherical *H. lacustris* cells with strong red autofluorescence compared to low-fluorescence elongated *E. gracilis* cells. Results yielded a sorting purity of 90.1% and a yield of 51.4%. A region of clumped of *E. gracilis* cells in the unsort channel presented a challenge to accurately count and an error was added to the cell count to account for this. Additional images of the sorting results are shown in Supplementary Fig. 4.

These label-free demonstrations firmly show the advantages of RACS over FACS. Although starch and paramylon particles can be quite large (~2 μm) and can make up a significant portion of cell mass, their tightly bound molecular structures make them difficult to stain with traditional fluorescent dyes and IgG-based antibodies, rendering them largely inaccessible to FACS machines. Two recent paramylon-targeted studies have made use of much smaller peptide^44^ and ssDNA-based^45^ aptamers for paramylon staining, but these are not easily available and require cell permeabilization. In the case of starch, one previous demonstration employed a negative detection scheme, that is, staining the cytosol and attributing regions of reduced fluorescence intensity to the presence of starch granules^46^. This neatly emphasizes the benefits of coherent RACS as demonstrated here: cell sorting based on the direct measurement of cellular contents and simple, label-free sample preparation, saving experimental time and cost.

## Discussion

In this article, we have demonstrated label-free broadband RACS at a record high-throughput of ~50 eps by combining a broadband fingerprint-region FT-CARS spectrometer, a dual-membrane push-pull sorter, and a real-time spectral processor, allowing real-time decision-making based on the cellular biochemical signal. Our RACS machine accomplished a two orders of magnitude increase in throughput for broadband RACS, effectively reducing the time necessary to sort a sample of 10,000 cells from about 5 hours in the fastest-reported broadband RACS machine to date^22^ to about 3 minutes. This increase in throughput is essential for the applicability of RACS to time-sensitive protocols, protocols requiring a large number of cells sorted, and for the enrichment of rare cells. To demonstrate the diverse applicability of our RACS machine, we have shown label-free sorting of plastic beads and of microalgae containing the carotenoid astaxanthin and the polysaccharides paramylon and starch, ranging in sizes from 3 to 50 μm. The label-free nature and background-free signal of the FT-CARS measurement in our RACS machine allow sorting cells without staining and in their original growth media, simplifying experimental workflow significantly compared to FACS and even spontaneous RACS. Our results indicate that the current fingerprint-region RACS system holds promise for microbiological, environmental, and bioengineering applications.

Although these results constitute a significant advancement in broadband RACS, our demonstration was limited to the detection of biomolecules with high concentration and/or large scattering cross-sections. For wider application of RACS, especially to mammalian cells and bacteria, further improvements in sensitivity, throughput, and apparatus cost are needed. First, sensitivity enhancements by heterodyne detection^47,48^, Sagnac interferometry^49^, polarization-selective measurement^50,51^, and quasi-dual comb laser systems^52^ have been reported for coherent Raman measurements, but not yet applied to flow cytometry or RACS. Sensitivity can also be enhanced by stepping out of the label-free regime and employing Raman tags^53^. Second, throughput can be enhanced by narrowing the sorting window and improving particle arrival spacing. Both have been demonstrated in a single device by utilizing pulsed-laser driven cavitation to sort cells at throughputs up to 6,000 eps with high purity in a microfluidic FACS device^54^. This device achieved its improvement to cell arrival spacing through the incorporation of an expansion chamber upstream of the measurement region, reducing double cell sort events. Third, the Ti:Sapphire pulsed laser and optics used in this system constitute a considerable expense that presents a challenge to widespread adoption. Therefore, a fiber laser-based FT-CARS implementation^55^ would lower the cost and size of the apparatus.

Several additional applications are possible with further development of our RACS technology. Fingerprint-region Raman image-activated cell sorting could be realized with spatial scanning of the FT-CARS beam focus, and the relaxed sampling requirements of compressive sensing^56^ could ameliorate throughput reductions that beam scanning may incur. Alternatively, multi-focus CARS using a microlens array^57^ could be implemented for flow-through imaging. Although not demonstrated here, RACS based on the detection of extracellular biomolecules should also be feasible via droplet microfluidics, which consists of micrometer water-in-oil droplets that sequester cells and their excretions. Isozaki *et al*.^58^ demonstrated sorting of cells in large (>100 pL) droplets at high throughput (>1,000 eps) using an on-chip dielectrophoretic array, which could be integrated into our RACS machine. Finally, more complicated spectral analysis methods such as principal component analysis or multivariate curve resolution could be implemented in our sorting algorithm to take further advantage of its broadband spectra.

## Materials and Methods

### FT-CARS Spectrometer

Our FT-CARS spectrometer consisted of a 80-MHz Ti:Sapphire femtosecond mode-locked laser (Coherent, Vitara T-HP) with a bandwidth of 88 nm and a center wavelength of 800 nm. Pulses from the laser were first directed through a pulse shaper (described below) to account for third-order pulse dispersion. Pulses then entered the pulse pair generator (PPG), which split each pulse into a pulse pair and rapidly scanned the optical delay between them. The PPG consisted of a rapid-scan Michelson interferometer, described in our previous work^24^, which split each incoming pulse using a halfwave plate and polarizing beam splitter. The pair traveled down opposite arms of the interferometer, with one arm rapidly scanning optical delay (tunable up to 1 mm, with 0.523 mm chosen for sorting experiments) using a 12-kHz resonant scanner (Cambridge Technology, CRS 12 KHz). Delay was accurately recorded by the interferogram from a 1064-nm narrowband laser aligned collinearly with the Ti:Sapphire laser light through the interferometer. Following the PPG, pulses were compressed by −4550 fs^2^ by a pair of chirped mirrors to remove second-order dispersion (Thorlabs, DCMP175), then passed through a long-pass filter (Thorlabs, FELH0750) to distinguish the anti-Stokes region from the laser bandwidth. A pair of lenses (Olympus, LCPLANN50X0.65IR) were used as condenser and objective at the sample. Anti-Stokes light produced at the sample was isolated by an optical short pass filter (Thorlabs, FESH0750) in the forward-scattering direction and detected by an avalanche photodiode (Thorlabs, APD120A/M). Acquisition triggering used the rising edge of forward scattering intensity from a 635 nm diode laser (Thorlabs, CPS635) focused through the same objective to a point in the flow channel upstream from the FT-CARS laser. The forward scattering laser was separated from the anti-Stokes signal by a polarizing beam splitter and focused onto a spatial mask. Cells passing through the laser focus in the flow channel modified the laser beam path, allowing light to pass the spatial mask to be detected by a photodiode (Thorlabs, PDA10AEC). For flow cytometry mode, data were digitized with a digitizer (AlazarTech, ATS9440) and post-processed with Igor Pro (Version 8, Wavemetrics) and Matlab (Version R2020b, Mathworks). The hardware and analytical methods for real-time spectral data collection, processing, and decision making for sorting mode are discussed in depth below.

### Real-time spectral processor

The time-domain FT-CARS interferogram, continuous-wave (CW) interferogram, and forward scattering signal were fed to the real-time spectral processor, which digitized them with a 16-bit digitizer module (National Instruments, NI5734) at a sampling rate of 120 MS/s and then processed these data with a field programmable gate array (National Instruments, PXIe-7972R) at a clock rate of 120 MHz. Each acquisition was triggered by the rising edge of the forward scattering signal, with signal recording starting at the first following 0 or π position of the resonant scanner cycle. The time-domain FT-CARS signal was stored in memory for every clock until the CW interferogram crossed zero voltage. At this zero-crossing, the stored FT-CARS signals were averaged to obtain one sample of a calibrated time-domain FT-CARS interferogram. This sampled the calibrated time-domain FT-CARS interferogram every 532 nm (1.77 fs in optical delay time). This sampling interval was short enough for our measurements, in which the maximum frequency of detected Raman peaks is about 1,600 cm^−1^, equivalent to a period of 20.8 fs. In a resonator half-period of 41.67 μs (equivalent to a single mechanical scan of optical delay by the interferometer), the signal acquisition ended when 1,024 calibrated FT-CARS samples were acquired, which happened when the pump-probe delay reached 0.546 mm (532 nm × 1,024). Then, the calibrated FT-CARS interferogram was Fourier transformed, which took 2,634 clocks (21 μs). The Fourier-transformed spectrum was then sent to the spectral averaging node as well as a windows computer via an NI PXI platform (National Instruments, PXIe-1071, PXIe-PCIe8381) for displaying and saving the obtained spectra. In the spectral averaging node, spectra were averaged then sent to the sort decision node. The number of spectra averaged was selected by users based on cell size, Raman signal intensity, and flow speed. The sort decision node received the spectral average for each event and calculated an event score based on the intensity ratio between two user-selected Raman shifts. This score was then compared to a user-set threshold value, with sorting actuated for scores above the threshold (or below, as toggled by the user in the software control). To actuate sorting when the target cell reached the correct point on the chip, the sort signal was sent to the on-chip sorter after waiting for a predetermined delay time (0 – 2,000 μs, user selected). In practice, the time between a particle entering the measurement position and the start of sort actuation was ~1.1 ms. To keep track of throughput across the experiment, the RACS machine recorded the total count of events at one-second intervals.

### Dual-membrane push-pull sorter

The dual-membrane push-pull sorter^27^ consisted of a three-layer, silicon-between-glass chip. Main channel geometry comprised a 200 μm × 200 μm cross-sectioned flow channel. Flowing cells were acoustically focused by a piezoelectric transducer glued to the bottom chip face and driven with a ~3.7 MHz, 50.0 V_pp_ sine wave. At the sorting region, the main channel split into three outlet branches, with reservoirs preceding the outlets on either side of the main channel (Fig. 2). For an unsort event, unmodified flow directed cells to exit through the central outlet to the chip’s unsort collection. For a sort event, the glass membranes of the reservoirs were deflected by piezoelectric actuators driven by a ramp voltage signal of 29 V at 64 μs rise time. This produced high-speed, local flow crossing the main flow channel, directing cells into either sort outlet, which joined down-chip at the sort collection.

### Cell sorting videos

To take videos of cell sorting, facilitate sample alignment, and monitor sorting performance in real time, a high-speed imaging system was integrated into the RACS machine. The system consisted of illumination from a green light-emitting diode (LED; Thorlabs, M530L4) counterpropagating along the FT-CARS beam path through the sample. Optical long pass filters (Edmund Optics, 86335) inserted the light into the beam path at the objective lens and picked it off at the condenser lens to be sent to a high-speed camera (Vision Research, Phantom Miro 4). A video was captured at 2,200 frames per second (fps) for analysis using manufacturer-provided software (Vision Research, Phantom Camera Control).

### Pulse shaper

The pulse shaper consisted of a dispersive prism (Thorlabs, PS859) coupled with a cylindrical concave mirror (Thorlabs, CCM400P01) and spatial light modulator (SLM; Santec, SLM200) in a retroreflective 4-f configuration with the SLM at the Fourier plane. A slight vertical misalignment in the pulse shaper 4-f allowed outgoing pulses to be selected by a mirror and sent to the PPG. Pulse shaping was performed using a genetic algorithm^59^ with fifty genes, a mutation factor of 0.06, and a pixel bin-size of 10 across the SLM. Fitness for early generations was determined by second harmonic signal generation from a beta barium borate (BBO) crystal placed at the sample position. Further pulse shaping was done on the microfluidic chip with fitness determined by the geometric mean (integration region 300 – 1,600 cm^−1^) of the four-wave mixing spectral intensity signal from water flowing in the channel.

### Preparation of polymer particles

Fluorescent PMMA (PolyAn, 5 μm pink) and PS (Spherotech, 7 μm yellow) microbeads were prepared in standard solutions, counted with Neubauer chambers, and mixed 1:1 to prepare the sorting samples. Sorting sample densities were at 1 × 10^5^ particles/mL for 31 eps sorting, and 5 × 10^4^ particles/mL for 13 eps sorting.

### Preparation of *E. gracilis* cells

*E. gracilis* (NIES-48) cells were cultured in ~8 mL of Koren-Hutner (KH) media pH 3.5 in T25 cell culture flasks laid flat to maximize gas exchange. Cells were incubated at 26°C with 14:10 h day:night cycles and 120 μmol photons/m^2^ s illumination. Cells grown under the day:night cycle made up the paramylon-poor cohort. Paramylon production was induced in the paramylon-rich cohort by seeding cells in fresh KH media at 5 × 10^4^ cells/mL. The flask was wrapped in tinfoil to block light and promote heterotrophy. Cells were incubated for four days before measurement. Both *E. gracilis* cohorts were fixed with 1% glutaraldehyde prior to sorting to facilitate counting (*E. gracilis* is an active swimmer and difficult to count when live). Fixed cell aliquots from the paramylon-rich and paramylon-poor cohorts were counted with Neubauer chambers, then mixed 1:1 to produce the sorting sample.

### Preparation of astaxanthin-rich *H. lacustris* cells

*H. lacustris* (NIES-4141) cells were cultured in 8 mL of AF6 media in a T25 cell culture flask. Cells were incubated at 26°C with 14:10 h day:night cycles and 120 μmol photons/m^2^ s. Astaxanthin-rich red cyst formation was complete after several weeks due to nutrient starvation. For sorting, *H. lacustris* red cysts were mixed 1:1 with low-paramylon *E. gracilis* cells.

### Preparation of *C. zofingiensis* cells

*C. zofingiensis* cells were grown in modified acetate (mAc) medium, prepared as AF6 medium with 0.4 g/L glucose, yeast extract, tryptone and sodium acetate. Prior to the experiment, cells were seeded at a concentration of 10^5^ cells/mL in either mAc medium (low glucose) or high-glucose mAc medium (20 g/L glucose). Cells were incubated at 26°C with 12:12 h day:night cycles and 120 μmol photons/m^2^ s for 7 days. On day 7, the low glucose sample presented a higher concentration (1.25 × 10^7^ cells/mL) of smaller cells (7.3 ± 3.5 μm in diameter), whereas the high glucose sample had a lower concentration (6.28 × 10^6^ cells/mL) of larger cells (9.3 ± 2.7 μm in diameter). To allow Lugol staining of the sorted cells, fixation, and permeation of the cells prior to sorting were required. Cells were fixed with ethanol (1:1). The ethanol-containing solution was washed and the sample was resuspended in 3 mL of ultrapure water and stored at 4°C for a week. After sorting, Lugol staining (5-10% I/KI) was added to the well containing the sorted cells to a final concentration of 2-4% I/KI. The bright-field images were counted manually using ImageJ and a point of intensity in the cell was used to classify the cell as starch-rich (dark cells) or non-starch-rich (light cells).

### Counting sorting results

Polymer microbeads and cells from sort and unsort collections were collected in slide-glass-bottom well dishes (Matsunami, D141400). These were centrifuged (5 minutes at 500 g) and then imaged with a Nikon TI eclipse fluorescence microscope (Nikon Instruments Inc.) equipped with a 20x objective lens (Nikon Instruments Inc., S Plan Fluor ELWD 20x) and a chlorophyll filter cube (Chroma Technology Corporation, 19010-AT-FM 1-43/Chlorophyll Longpass) and controlled with NIS-elements software (Nikon Instruments Inc.). Cells were counted using Gnu Image Manipulation Program to manually tag cells in the images and ImageJ was used for automated counting of the tags.

### Sorting purity vs throughput

Sorting performance for high-throughput cell sorters faces an upper bound determined by the probability of a sort event containing more than one cell. We define a “sort window” as the time required to mechanically execute a sort action. If the arrival times of two cells at the measurement point are within the same sort window and one is a target cell, the additional cell can be pushed into the sort channel along with the target. If this additional cell is a non-target cell, it reduces purity. As throughput increases, the arrival interval between cells shortens (following a Poisson distribution), creating a natural inverse relation between throughput and sorting purity. We simulated the maximum Poisson-allowed purities for our sort window (4.1 ms) at throughputs up to 60 eps, following the method established in Nitta *et al.*^12^. The ratio of false-positive and true-positive sort events can be modified by the sample ratio of target and non-target cells, further modifying the theoretical maximum sorting purity at a given throughput.

### Calculation of purity, yield, and enrichment

Sorting results were calculated from microbead or cell counts C as

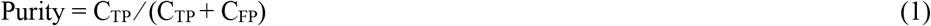

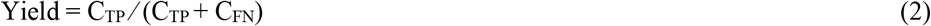

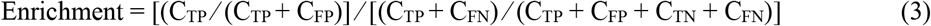

where counts were defined as

C_TP_ ≡ the number of target cells in the sort collection (true positive events)
C_FP_ ≡ the number of non-target cells in the sort collection (false positive events)
C_TN_ ≡ the number of non-target cells in the unsort collection (true negative events)
C_FN_ ≡ the number of target cells in the unsort collection (false negative events)

## Data availability

The datasets generated during and/or analyzed during the current study are available from the corresponding author upon reasonable request.

## Code availability

All codes used for analysis of this study are available from the corresponding authors upon reasonable request.

## Acknowledgments

This work was supported by KISTEC, JST PRESTO (JPMJPR1878), JSPS Core-to-Core Program, JSPS Grant-in-Aid for Young Scientists (20K15227), Grant-in-Aid for JSPS Fellows (19F19805), Nakatani Foundation, White Rock Foundation, Ogasawara Foundation, and Kurita Foundation.

## Author contributions

K.G. conceived the idea of RACS. K.H. and K.G. designed the system. M.L., J.G.P., and K.H. developed the optical setup. K.H. developed the code of the sorting program. M.L., J.G.P., J.W.P., A.I., and K.H. performed the sorting experiments and analyzed the data. M.L., J.G.P., J.W.P., A.I., K.H., and K.G. wrote the manuscript. K.G. supervised the work. All authors discussed the results.

## Author Information

Reprints and permissions information are available at www.nature.com/reprints. Correspondence should be addressed to K.H. and K. G.

