## Supplementary Material for "High-throughput Raman-activated cell sorting in the fingerprint region"

for

**This PDF file includes:**

Supplementary Figs. 1 to 4

**Other Supplementary Materials for this manuscript include the following:**

Supplementary Movies 1 to 2

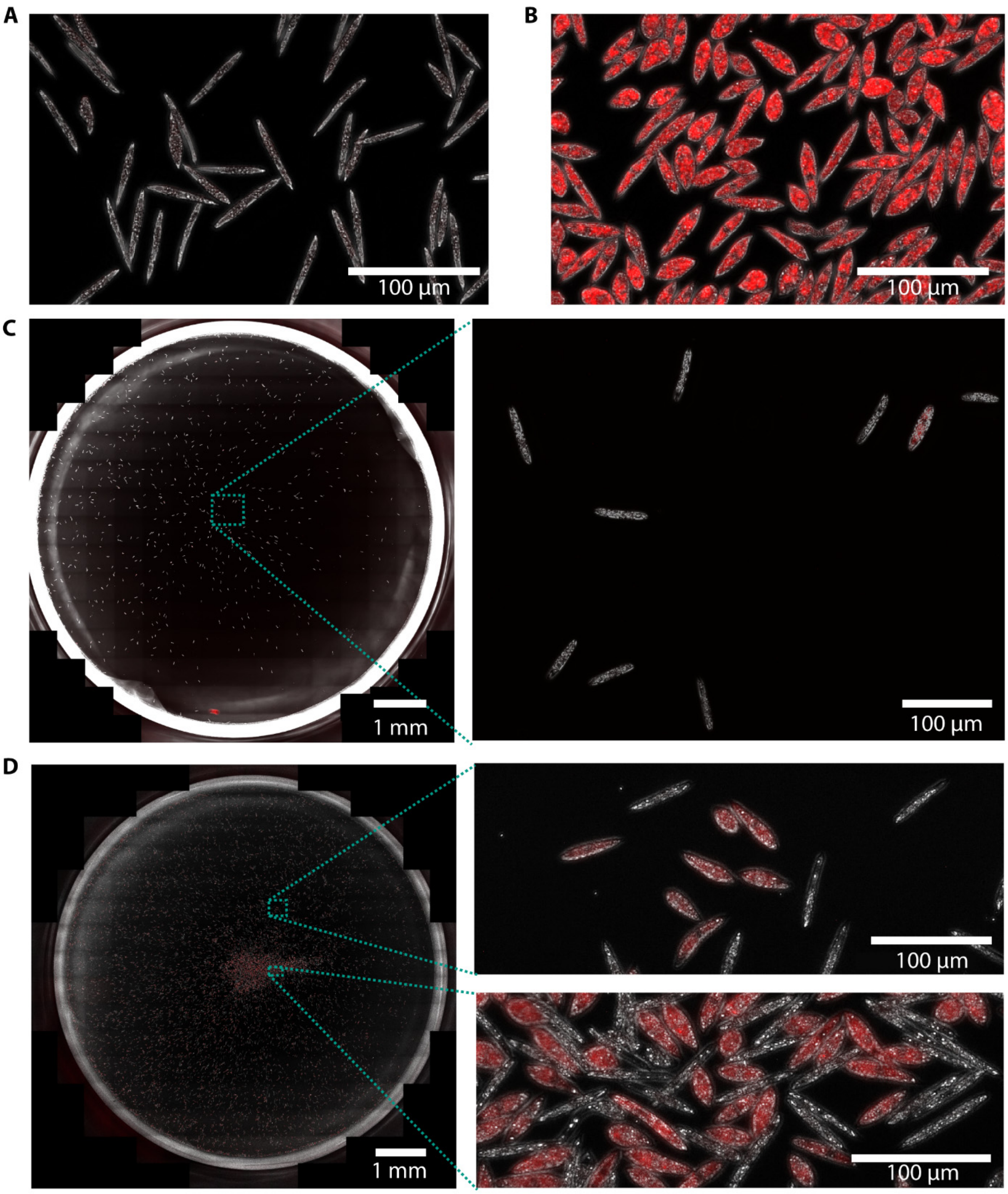


**Supplementary Fig. 1 | Bright-field and fluorescence images from sorting *E. gracilis* cells for paramylon content.** (**a**) *E. gracilis* sample from cells grown under heterotrophic (paramylon-promoting) conditions. These cells lack chlorophyll autofluorescence, shown in the red channel. (**b**) Cell sample grown under mixotrophic (control) conditions. These cells have strong chlorophyll autofluorescence, shown in the red channel. Images of the (**c**) sort collection and (**d**) unsort collection after sorting a mixed sample of heterotrophic and mixotrophic cells.

**A**

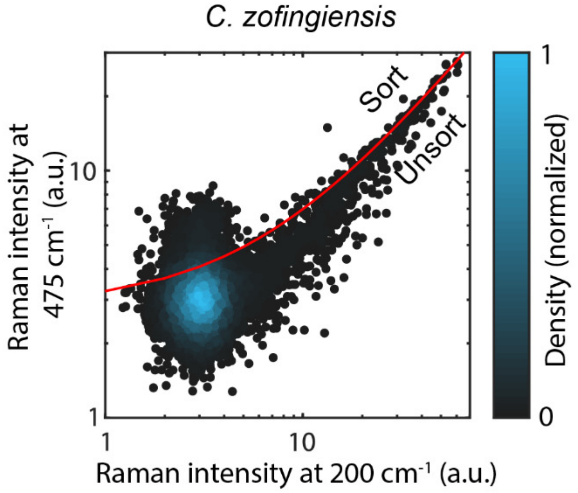


**B**

| | Sort | Unsort | Total |
|---|---|---|---|
| High starch | 435 | 528 | 2773 (6%) |
| Low starch | 869 | 13948 | 3461 (94%) |

| | |
|---|---|
| Throughput | 40 ± 10 eps |
| Purity | 33.4% |
| Yield | 45.2% |
| Enrichment | 5.4× |

**Supplementary Fig. 2 | Sorting of a low target ratio sample.** (**a**) Scatter plot from *C. zofingiensis* mixed phenotype cell sorting to select starch-rich cells, with (**b**) counted results.

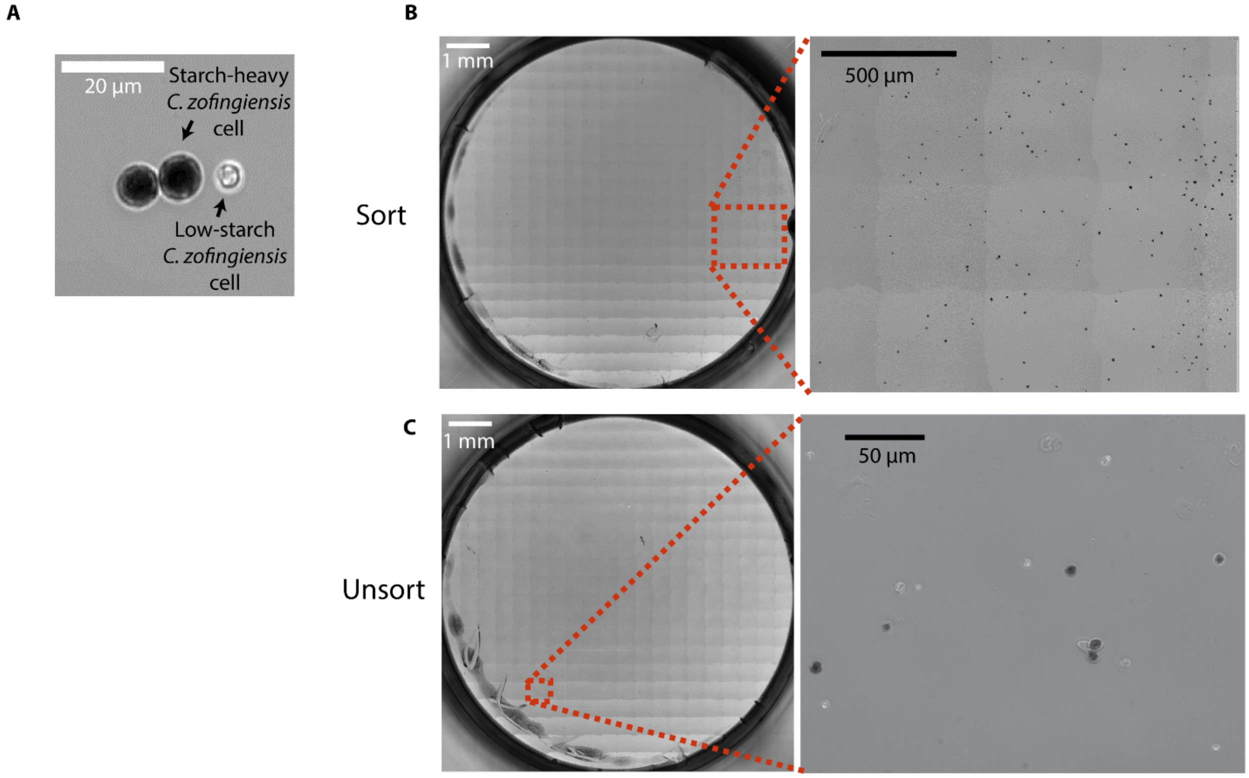


**Supplementary Fig. 3 | Bright-field images of Lugol stained *C. zofingiensis* cells of a 1:1 mix sample following sorting.** (**a**) Example of bright-field images cells following Lugol staining, showing two starch-heavy cells with dark staining and a low-starch cell, appearing white. (**b**) Bright-field images of the well and zoom in for the sort collection, showing an enrichment of the target starch-heavy dark cells (79.8%). (**c**) Bright-field images of the well and zoom in for the unsort collection, showing lower percentage of target cells (33%).

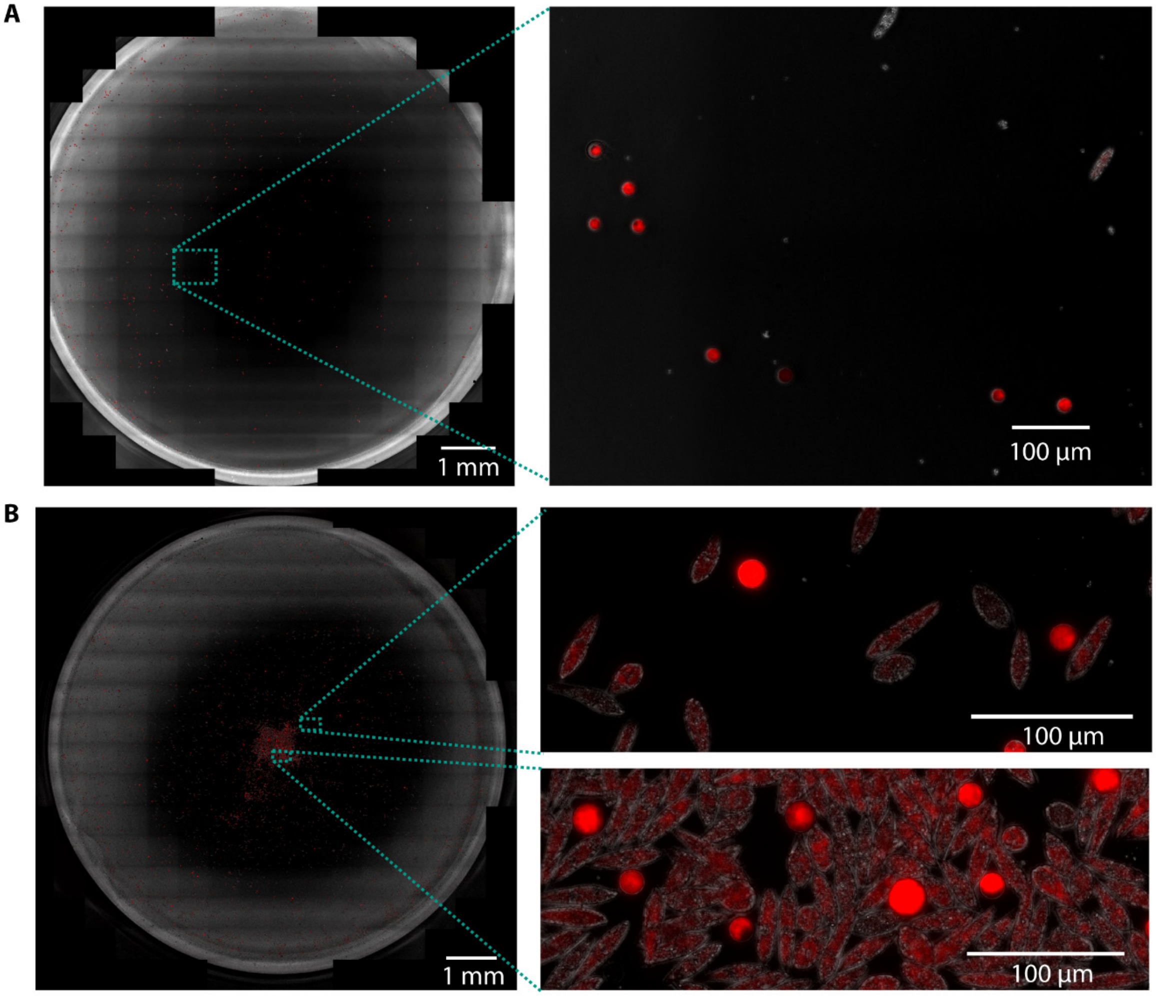


**Supplementary Fig. 4 | Bright-field and fluorescence images from sorting *H. lacustris* and *E. gracilis* cells for astaxanthin content.** (**a**) Sort collection. (**b**) Unsort collection. *H. lacustris* cells appear as spherical cells with strong autofluorescence from chlorophyll in the red channel. *E. gracilis* cells appear as rod shaped cells with moderate autofluorescence.
